# Dual-Targeted Microbubbles Promote Apoptosis of M1 Macrophages by Inhibiting Telomerase Activity for Atherosclerosis Therapy

**DOI:** 10.1101/2024.08.01.606265

**Authors:** Wei Zeng, Zhengan Huang, Yalan Huang, Kaifen Xiong, Yuanyuan Sheng, Xiaoxuan Lin, Xiaofang Zhong, Jiayu Ye, Yanbin Guo, Gulzira Arkin, Jinfeng Xu, Hongwen Fei, Yingying Liu

## Abstract

It is known that M1 macrophages are the dominant phenotype in progressive atherosclerosis (AS). Studies have shown that telomerase in macrophages can be activated during the atherosclerosis formation, but how to regulate the telomerase activity of M1 macrophages is the focus of treatment of AS. Herein, we report a dual-targeted microbubbles delivery system encapsulating the telomerase inhibitor BIBR1532 (Ab-MMB_1532_), designed for precise targeted therapy of AS. The Ab-MMB_1532_ exhibits excellent biocompatibility and remarkable targeting capability towards M1 macrophages with its targeting advantage notably accentuated under high shear forces. Furthermore, it significantly downregulates telomerase activity. Metabolomics analysis indicates that the decrease in telomerase activity upregulates the expression of aspartic acid, a key protein in the Caspase pathway of apotosis. In vivo, Ab-MMB_1532_ efficiently targeted and accumulated within AS lesions, leading to significantly delay of the AS progression by inhibiting telomerase activity and promoting apoptosis of M1 macrophages after 4-week treatment. In summary, Ab-MMB_1532_ provides a new approach for potentially safer and more effective treatment of AS in the future.

## INTRODUCTION

Atherosclerosis (AS), a chronic inflammatory artery disease, characterized by the accumulation of lipid and fibrous elements deposition underneath the inner wall of vessel, is a major cause of cardiovascular death, taking an estimated 17.9 million lives each year^1,2^. The growth of atherosclerotic lesions or the rupture of ‘vulnerable’ plaques can lead to acute myocardial infarction, stroke and sudden cardiac death^3,4^. In that case, early diagnosis and intervention of AS are vital for slowing the disease and saving more lives. Currently, lowering lipid levels, controlling hypertension, antiplatelet therapy and preventing blood clot formation have become routine clinical treatments for AS. However, the incidence and mortality of AS remain significantly high. Moreover, severe AS may require percutaneous and surgical management, albeit often effective, depends highly on expensive and invasive technology^5^. Therefore, it is urgently needed to develop novel therapeutics strategies for atherosclerosis.

There is strong evidence that macrophages are a key player in atherosclerotic disease progression and regression^6^. In AS, macrophage dysregulation underlies the disease pathogenesis^7^. During the initial phases of AS, the differentiation of monocytes into macrophages is drived by macrophage colony-stimulating factors and other differentiation factors. During the development of AS, a large amount of aggregated macrophages, particularly M1 macrophages, secreting inflammatory cytokines (TNF-α, IL-1β and IL-6) and leading to foam cell formation^8^. These evidences indicate that regulating the inflammatory state and function of macrophages, especially M1 macrophage, raises hope for AS regression^9^. One question is how we can target M1 macrophages as a therapeutic strategy for anti-AS.

Nanotechnology is particularly advantageous for addressing these challenges^10^. Recent advances have shown the potential of targeting drug delivery system based on ultrasound microbubbles (MBs) for anti-AS^11–14^. However, considerable challenges remain in targeting efficiency and controllable drug release at the lesion sites caused by the shear forces of blood flow^15,16^. Fortunately, the emerging magnetic ultrasound microbubble (MMB) navigation technology has the ability to alter the axial distribution characteristics of microbubbles within blood vessels. Under the influence of an external magnetic field, MMB can overcome the shear forces, pushing MMB closer to the vessel wall, thereby increasing their opportunities for contact with target sites on the region of interest^17–19^.

Telomerase is a ribonucleoprotein complex that consists of several components including TERT, telomerase RNA component that serves as a template for adding telomere repeats^20^. It has been reported that the infiltration of macrophages during the progression of AS, has the capacity to reactivate telomerase, thereby inducing an up-regulation in TERT expression^21,22^. BIBR1532 is a synthetic nonnucleoside, noncompetitive small-molecule telomerase inhibitor that can inhibit telomerase activity by specifically binding to the active site of TERT ^23,24^.

Hence, based on the above background, we investigated a dual-targeted ultrasound microbubble delivery system loaded with BIBR1532 (Ab-MMB_1532_) for the targeted therapy of AS (as shown in the Schematic illustration). The results showed that Ab-MMB_1532_ could target and accumulate on M1 macrophages within AS plaques through an external magnetic field and CD86 antibodies, then the ultrasound cavitation effect would promote local release of BIBR1532 into M1 macrophages, resulting inhibition of telomerase activity and inducing their apoptosis. Correspondingly, metabolomics analysis has confirmed the decrease in sphingosine metabolism and the upregulation of aspartic acid. Finally, in vivo studies demonstrate that Ab-MMB_1532_ exhibits optimal efficacy against AS.

**Schematic illustration.**
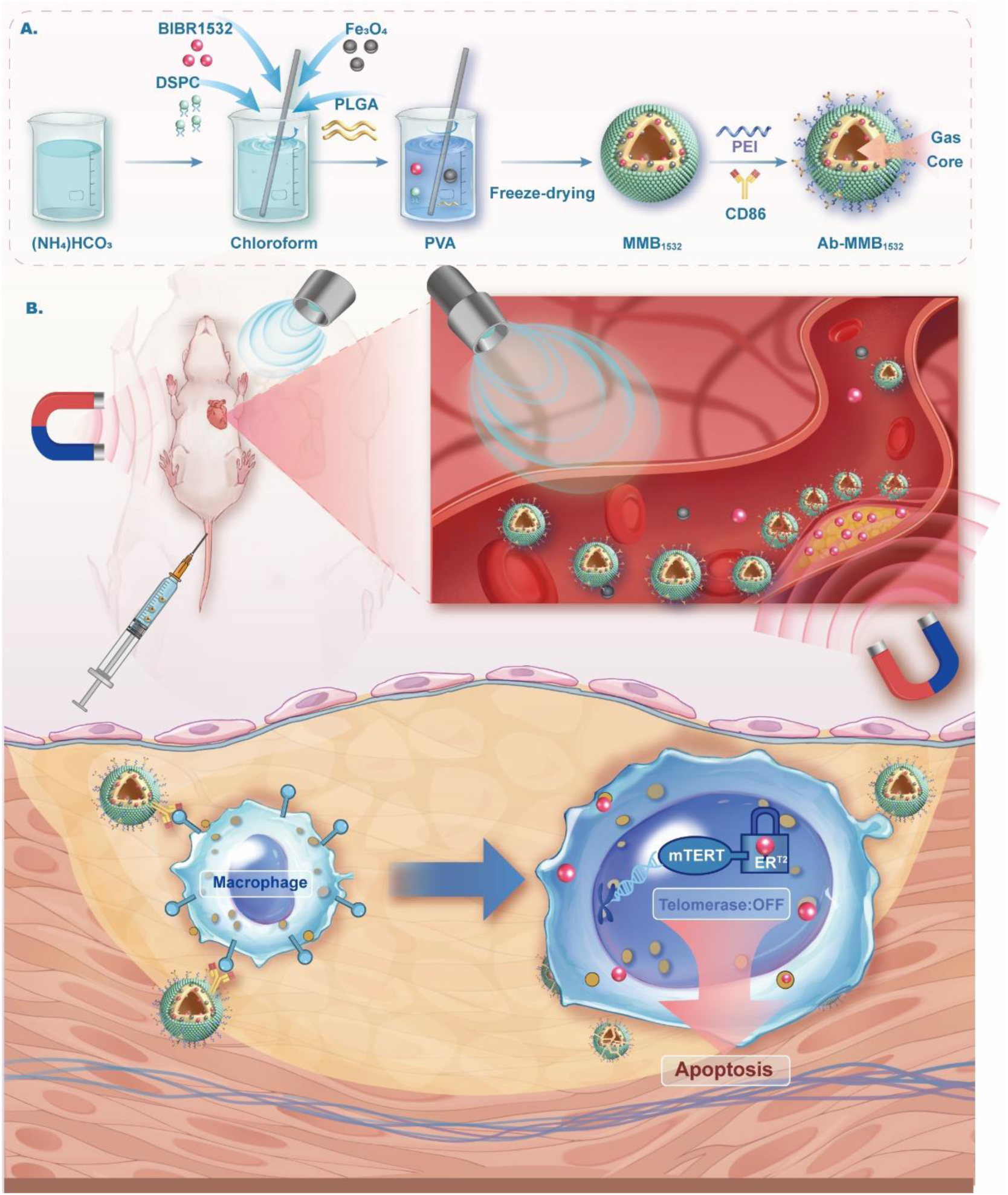
The Ab-MMB_1532_ fabrication, targeting, and its anti-AS properties.

## MATERIALS AND METHODS

### Antibodies and reagents

Antibodies against iNOS, CD86, Caspase-3 and Cleaved Caspase-3 were purchased from Cell Signaling Technology (Boston, USA). Antibodies against TERT were purchased from Proteintech Group (Wuhan, China). Antibodies against β-actin were purchased from Affinity Biosciences (California, USA). Anti-rabbit HRP-conjugated IgG secondary antibody, lipopolysaccharide (LPS), Ammonium Bicarbonate ((NH_4_)HCO_3_), polyvinyl alcohol (PVA) were purchased from Sigma-Aldrich (St. Louis, USA). Dulbecco’s modified Eagle’s Medium (DMEM), fetal bovine serum (FBS) were purchased from Invitrogen (CA, USA). 1,2-distearoyl-snglycero-3-phosphocholine (DSPC) were obtained from Avanti Polar Lipids (AL, USA). BIBR1532 were bought from MCE (New Jersey, USA). 100× penicillin-streptomycin solution, 1,1’-dioctadecyl-3,3’,3’-tetramethylindocarbocyanine perchlorate (DiI), 2-(4-Amidinophenyl)-6-indolecarbamidine dihydrochloride (DAPI) were purchased from Beyotime Biotechnology (Shanghai, China). Poly (DL-lactide-co-glycolide) (PLGA, 50:50, MW = 30 000) were purchased from Jinan Daigang Biological Material (Shandong, China). Oleic acid-modified Fe_3_O_4_ nanoparticles were bought from Nanjing Nanoeast Biotech (Nanjing, China).

### Synthesis of MB_1532_, Ab-MB_1532_, MMB_1532_ and Ab-MMB_1532_

The MB_1532_ and MMB_1532_ were fabricated by a modified double-emulsion-solvent-evaporation method^25^. Briefly, 50 mg of PLGA, 2.5 mg DSPC and a certain amount of BIBR1532 dissolved in 1 mL chloroform. Then 0.2 mL freshly prepared (NH_4_)HCO_3_ solution (60 mg/mL) was added to the above oil phase. The mixture was emulsified using an ultrasonic probe in an ice bath for 2 min, with a pulse mode of working 3 s and resting 3 s. Subsequently, the initial emulsified solution was slowly added into 5 ml PVA (4%, w/v), and then homogenized for 5 min at 6000 rpm. Incorporate 10 mL of pre-cold deionized water into above obtained emulsion, and stir magnetically for 4 hours at RT to evaporate the organic solvent. After that, Collect the precipitate by centrifugation at 6000 rpm and 4°C for 10 minutes, followed by three washes with deionized water. Finally, resuspend the precipitate in 0.5 mL of deionized water, followed by vacuum freeze-drying for 24 hours to obtain MB_1532_, which stored at 4°C for use. Following the same method above, 3 mg oleic acid-modified Fe_3_O_4_ nanoparticles or 2μL DiI was added into the oil phase to obtain the production of distinct entities: magnetically responsive MMB_1532_ and DiI-MB_1532_ / MMB_1532_ with red fluorescent labeling.

The MB_1532_ and MMB_1532_ were further coated with PEI to impart a positive charge, and then 20 μL of CD86 antibodies were added to obtain Ab-MB_1532_ and Ab-MMB_1532_ respectively, through electrostatic adsorption.

### Characterization

The morphology of MB_1532_ and MMB_1532_ were analyzed by scanning electron microscope SEM (Hitachi, Japan) and transmission electron microscope TEM (JEOL, Japan). The particle size and zeta potential of MB_1532_ and MMB_1532_ were determined by Zetasizer NanoZS instrument (Malvern, UK). Vibrating sample magnetometer (VSM, Quantum Design, USA) was used to obtain the magnetization curve of MMB_1532_. To demonstrate whether Ab-MB_1532_ and Ab-MMB_1532_ successfully loaded CD86, fluorescence secondary antibodies (Dylight 649, Goat anti-mouse IgG) at the recommended concentration were incubated with MB_1532_ and MMB_1532_ for 10 minutes under protecting from the light at 4°C, followed by observation under a confocal laser scanning microscope (Leica TCS SP8, Germany). In order to observe the magnetic response performance of MMB_1532_, a rectangular magnet (25 × 10 × 5 mm, 1.2T) was positioned near a glass vial containing an MMB_1532_ aqueous solution, and a camera was employed to record the process of magnet attraction. To test the stability of MB_1532_, the zeta potentials of MB_1532_ were measured at 0 h, 12 h, 24 h, 2 d, 3 d, 4 d, 5 d, 6 d, 7 d, and 14 d after preparation.

### Cell culture and M1 macrophage polarization stimulation

Raw264.7 murine macrophage cells were purchased from ATCC (cat #TIB-71), and cultured at 37℃ with 5% CO_2_ in DMEM supplemented with 10% FBS and 1% penicillin-streptomycin. When reached 80% confluence, adherent raw264.7 cells were detached via scraping and then seeded in 6-well plates at a concentration of 3 × 10^5^ cells per well.

For M1 macrophage polarization stimulation, 100 ng/mL of LPS was added to the cell suspension mentioned above and mixed thoroughly, then seeded into a 6-well plate at a density of 3 × 10^5^ cells per well. The cells were incubated at 37°C in a CO_2_ incubator for 24 hours in preparation for the subsequent experiments.

### Cell viability assay and Flow cytometric analysis

Cell viability was assessed using Cell Counting Kit-8 (CCK-8; TargetMol). M1 macrophages were cultured with varying concentrations of BIBR1532 or microbubbles for 24 h. Following the kit instructions, cell viability was determined using the standard protocol. Experiments were conducted in triplicate.

Cell apoptosis was evaluated by flow cytometry (FC). Briefly, following treatment of M1 macrophages with various concentrations of BIBR1532, cells were gently washed twice with pre-chilled PBS for a duration of 3 minutes each. Subsequently, the cells were harvested by cell scraping and centrifuged at 1000 rpm for 4 minutes at 4°C, and the supernatant was discarded. Cell staining was performed according to the instructions provided in the Annexin V-PE / 7-AAD (Servicebio, China) cell apoptosis detection kit, with the entire procedure conducted in the absence of light and on ice. Cell apoptosis was assessed using a CytoFLEX flow cytometer (Beckman, USA) with FlowJo analysis software version 10.9.

### Telomerase activity assay and Protein Silver Stain Kit

Telomerase activity was measured with the Fluorescent RT-qPCR assay kit (Key-GEN BioTECH, China). PCR products were separated by electrophoresis on a 10% nondenaturing polyacrylamide gel (Beyotime, China), visualized by Protein Silver Stain Kit (Servicebio, China) staining.

### Senescence-associated β-galactosidase detection

Cells were subjected to senescence-associated β-galactosidase (SA-β-Gal) staining using a Beyotime Senescence Cell Staining Kit (Beyotime, China). After a PBS washing, cells were fixed for 15 minutes with fixing solution, followed by overnight incubation in staining solution at 37°C. Senescent cells were identified by green staining.

### Metabolomics analysis

For metabolomics analysis, M1 macrophages were treated with or without BIBR1532 for 24 hours. Samples (three biological replicates) were extracted with pre-cooled methanol : acetonitrile : H_2_O (2:2:1, v/v/v) and vortexed. Extracts were injected into the UPLC-MS/MS system for analysis. Raw data were converted to .mzXML format using ProteoWizard, followed by peak alignment, retention time correction, and peak area extraction using XCMS software. Data extracted by XCMS underwent metabolite structural identification and preprocessing, followed by quality assessment. Quality control (QC) samples were used for batch standardization. Differential analysis (t-test or ANOVA) was performed to determine p-values, adjusted p-values, fold change (FC), coefficient of variation (CV), and differential metabolites (P-value < 0.05 and FC < 0.67 or FC > 1.5). Pathway analysis was conducted using MetaboAnalyst 5.0 metabolomics software. Drug dose: BIBR1532, 30 μM.

### Optimal ultrasound therapy acoustic parameters in vitro

M1 macrophage cells in logarithmic growth phase were seeded into a 24-well plate at a density of 5 × 10^4^ cells per well, with 500 μl of culture medium in each well, and incubated at 37°C in a CO_2_ incubator for 24 hours. Different ultrasound parameters (intensity of 0.5, 0.8, 1.0, 1.2, 1.5, and 1.8 W/cm^2^; duty cycle of 10%, 20%, 30%, 40%, 50%, and 60%; treatment time of 10 s, 20 s, 30 s, 40 s, 50 s, and 60 s; Sonitron GTS Sonoporator, Japan), with untreated cells serving as a control. After another 24 h incubation, the cells were washed for three times using PBS, followed by staining by Live / Dead Cell Kit (KeyGEN, China). Briefly, the cells were incubated with Calcein-AM / PI solution (4 μM for each fluorescent probe in PBS) at room temperature for 45 minutes, followed by three washes with PBS, and imaging was performed under an inverted fluorescence microscope (Laica DMi8, Germany). The excitation wavelengths for Calcein-AM and PI were 488 nm and 543 nm, respectively, with corresponding emission bands collected in different wavelength ranges, specifically 510-540 nm (green) and 570-620 nm (red).

### CD86 antibodies and magnet-mediated BIBR1532 targeting transport in vitro

In order to assess the dual-targeted mediated biological effects, a modified inverted method was used, as we described previously^26^. In brief, this section of the experiment was divided into 6 groups (n = 4/group): the control group, free BIBR1532 group (30 μM), and 4 microbubble groups (MB_1532_, Ab-MB_1532_, MMB_1532_ and Ab-MMB_1532_). Microbubble suspensions were prepared by mixing fresh complete DMEM medium with a microbubble concentration of 2.5 × 10^7^ particles per mL and a BIBR1532 content of 30 μM.

M1 macrophages were then seeded in a 24-well plate at a density of 5 × 10^4^ cells per well. When the cells reached approximately 70% confluence, the medium was removed, and 3 ml of the prepared microbubble suspension was added to each well. Microseal ’B’ Adhesive Seals (Bio-Rad, USA) was used to seal the cell culture plate, and magnets were placed at the bottom of plates in all groups, followed by mildly rotation and removement of magnets for 10 minutes. After that, the supernatant was removed, gently rinsed twice with PBS to remove free unbound microbubbles. Then, 500 µl of fresh culture medium was added to each well, followed by a 40 s (1.2 W/cm^2^ of intensity, 30% of duty cycle) ultrasound treatment, and further overnight incubation. Protein expression was analyzed through Western blot (WB), and telomerase activity was assessed using a telomerase activity assay kit.

### Parallel plate flow chamber assay

A home-made parallel plate flow chamber assay was used to analysis the magnetic & antibody targeting attachment, as we described previously^27^. In brief, 1 mL of the preprepared Dil-labeled microbubble solution (MB_1532_, Ab-MB_1532_, MMB_1532_ and Ab-MMB_1532_) at a concentration of 2.5 ×10^7^ particles per mL, was introduced into a vacuum parallel plate flow chamber (Glycotech 31-001, USA) by a stepping motor (Yuhui, China) via a capillary tube (*r* = 15 mm) at shear forces ranging from 6 to 48 dyn/cm². Throughout the entire experimental procedure, a rectangular magnet was positioned above the chamber to capture the magnetic microbubbles (MMB_1532_ and Ab-MMB_1532_), while M1 macrophages were prepared below the chamber. Subsequently, the average red fluorescence intensity surrounding individual M1 macrophage was observed under the EVOS FL auto cell imaging system (Thermo Fisher Scientific, USA), allowing for the assessment of their targeting capability.

### Targeting to AS plaques in vivo

ApoE^-/-^ mice (6-week-old) were purchased from Vital River Laboratory Animal Technology (Beijing, China). All animal experiments were approved by the Institutional Animal Care and Use Committee of Jinan University (No: 20210225-18). In order to induce the formation of AS, all apoe-/- mice were fed with a high-fat diet (HFD, including 41% fat and 17% protein) for 8 weeks prior to the experiment below. Saline, MMB_1532_, AB-MB_1532_ and Ab-MMB_1532_ were administered with a dose of 2 mg/kg via the tail vein, followed by a magnet placed on the trunk side of the heart level for 5 min in each group of mice. After 24 hours, euthanasia was performed on the mice, and perfusion with pre-cooled PBS containing 4% paraformaldehyde was conducted to remove unbound MBs. The separated aorta and major organs were imaged and quantified for fluorescence using the Xenogen IVIS 200 system.

### Animals and treatment protocol

After 8 weeks of HFD feeding, randomly divided apoe^-/-^ mice into 4 groups (n = 6/group): (1) saline group; (2) MMB_1532_ group (0.04 ng/kg BIBR1532); (3) Ab-MB_1532_ group (0.04 ng/kg BIBR1532); (4) Ab-MMB_1532_ group (0.04 ng/kg BIBR1532). All groups received weekly treatments for a consecutive duration of four weeks with 2-minute ultrasound irradiation (1.5 W/cm^2^, 30% duty cycle) at the anterior region of the mouse heart. Besides, for all groups, a magnet was initially placed on one side of the mouse torso for a 5-minute period, after which the magnet was removed before initiating ultrasound irradiation. All were fed high-fat diet during the treatment.

After 4-week period of treatment, all mice were euthanized, and the aorta and heart were carefully dissected, with whole blood collected for hematological analysis. The aorta was longitudinally dissected for Oil Red O (ORO) staining to observe AS plaques macroscopically. After cardiac perfusion, multiple consecutive 8μm-thick sections were prepared for H&E staining, ORO staining, and IF staining. Be-sides, H&E staining was performed on major organs (heart, liver, spleen, lung, and kidney) to assess the bio-safety of the therapeutic agents.

### RT-qPCR

Following the manufacturer’s protocol, total RNA was extracted from the cells using TRIzol^®^ reagent (Invitrogen, USA). The first-strand cDNA synthesis was per-formed using PrimeScipt^TM^ RT Master Mix (Takara, China). The cDNA and primers were mixed with TB Green^®^ Premix Ex Taq^TM^ II (Takara, China), and RT-qPCR was conducted on the StepOnePlus Real-Time PCR System (Thermo Fisher Scientific, USA) using the following thermal cycling program: 95°C for 30 s, followed by 40 cycles of 95°C for 5 s, 60°C for 30 s. β-actin was used as the endogenous reference gene, and relative gene expression was quantified using the 2^-ΔΔCq^ method^28^. The primers employed in this study were detailed in Table S2

### Western blot

The cells, following the treatment above, were collected and subjected to cell lysis using RIPA buffer (Beyotime, China). Subsequently, the protein concentration was determined using a BCA assay kit (Sparkjade, China). Sodium dodecyl sulfate-polyacrylamide gel electrophoresis (SDS-PAGE) and Western blot (WB) were carried out in accordance with standard procedures.

### Immunofluorescence staining of macrophages

Identification of M1 macrophages was carried out using IF staining. Cryosections or cell smears, following treatment, were fixed with 4% paraformaldehyde at room temperature for 30 minutes and then blocked with 10% goat serum for 1.5 hours. The primary antibody used were F4/80 antibody (1: 100, Santa Cruz, USA) and CD86 antibody (1: 200, Proteintech, China), and the secondary antibody used were anti mouse IgG Dylight 488 conjugated antibody (1: 200, Abbkine, USA) binding to F4/80 antibody and anti-rabbit IgG Dylight 649 conjugated antibody (1: 200, Abbkine, USA) binding to CD86 antibody. Cell nuclei were stained with DAPI. Glass slides were imaged using a confocal microscope, and image quantification was performed using ImageJ software.

### Statistical analysis

Statistical analysis was performed using SPSS 17.0 software (SPSS Inc., USA). Da-ta from in vitro experiments were presented as mean ± standard error (SE), and assessed using a 2-tailed Student’s t-test. A significance level of *p* < 0.05 was considered to be statistically significant.

## RESULTS AND DISCUSSION

Synthesis and characterization of MB_1532_, Ab-MB_1532_, MMB_1532_ and Ab-MMB_1532_ The Ab-MMB_1532_ was successfully prepared using the water/oil/water (W_1_/O/W_2_) double emulsion method and electrostatic adsorption (Figure 1A)^25^. The SEM image showed that MB_1532_ and MMB_1532_ exhibited a hollow spherical morphology, which were especially obviously in TEM. Furthermore, in comparison to MB_1532_, numerous black Fe_3_O_4_ nanoparticles were observed encapsulated within the bubble shell of Ab-MMB_1532_ (Figure 1B). The average particle sizes of MB_1532_ and MMB_1532_ measured by dynamic light scattering (DLS) were 1398 ± 17.72 nm and 2291 ± 157.09 nm, respectively, with their respective polydispersity indices (PDI) stood at 0.26 ± 0.07 for MB_1532_ and 0.23 ± 0.10 for MMB_1532_ (Figure 1B, Table S1). Before PEI modification, the zeta potential of MB_1532_ and MMB_1532_ were -18.33 ± 0.86mV and -21.23 ± 1.46mV, respectively. However, following positive charge modification, the potentials of MB_1532_ and MMB_1532_ significantly increased to 14.43 ± 1.34 mV and 20.27 ± 1.12 mV, respectively. After incubation with CD86 antibodies, it’s potentials slightly decreased to 7.62 ± 0.36 mV and 9.77 ± 0.46 mV (Figure S1). In addition, during 14 days follow-up in PBS solution, the zeta potential of MB_1532_ remained virtually unchanged (*p* > 0.05), suggesting that the MB_1532_ had remarkable stability and dispersion characteristics (Figure 1F, S2).

**Fig. 1.**
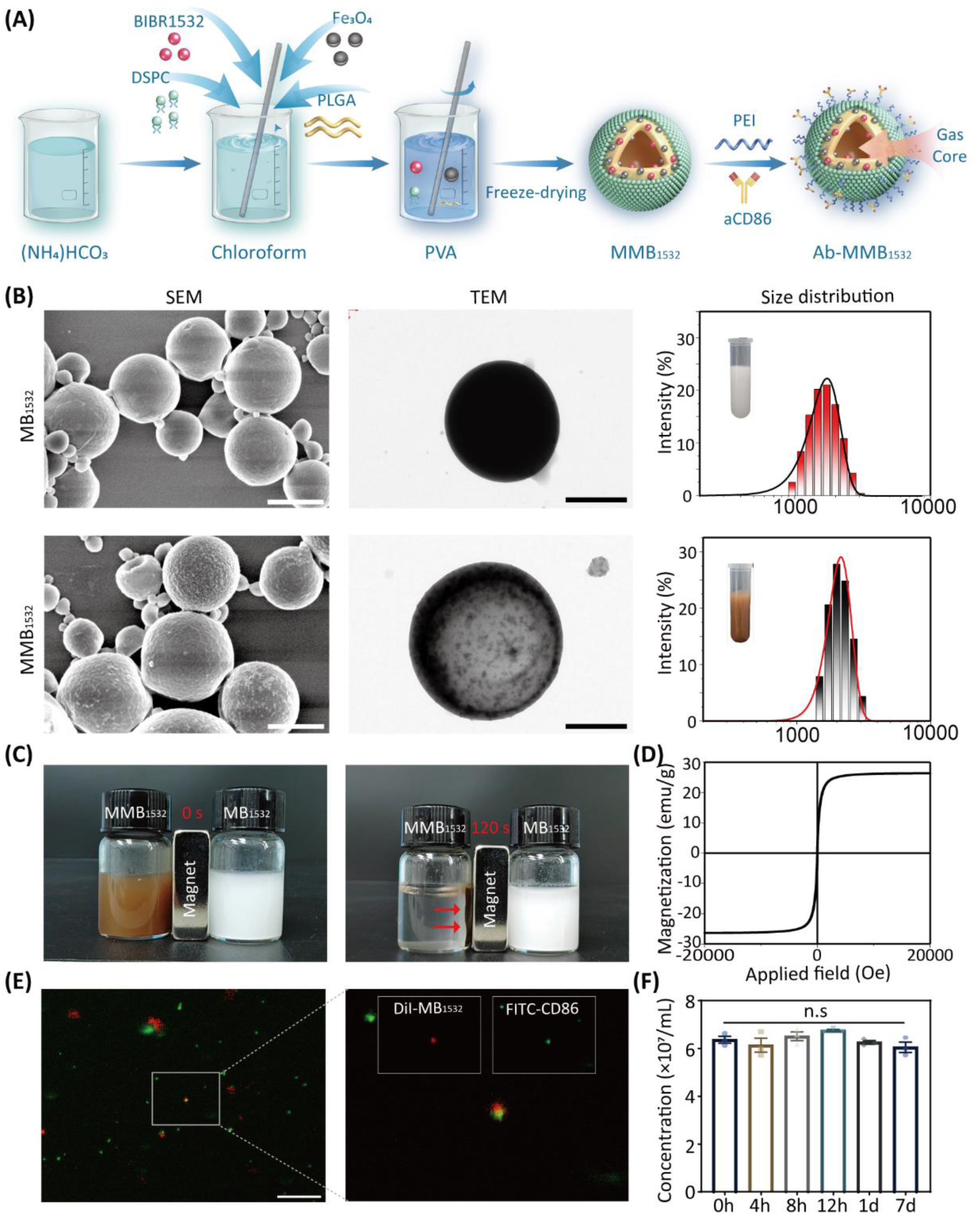
Fabrication and characterization of Ab-MMB_1532_. (**A**) Schematic illustration of the preparation of Ab-MMB_1532_. (**B**) SEM, TEM, and size distribution of MB_1532_ and MMB_1532_ respectively. Scale bar: 2 µm. (**C**) Comparative images of MMB_1532_ and MB_1532_ solutions before and after applying a magnetic field. (**D**) Field-dependent magnetization curve of MMB_1532_. (**E**) IF images of Ab-MB_1532_ show successful loading of CD86 antibodies (green) onto MB_1532_ (red). Scale bar: 25 µm. (**F**) Stability of the MB_1532_ over a 7d period at 37 °C.

Interestingly, the uniformly dispersion of MMB_1532_ in PBS was observed to be rapidly migrated towards the magnetic field’s orientation within a span of 120 seconds under the presence of a magnetic field (Figure 1C). Furthermore, it is worth noting that the field-dependent magnetization curve indicated that MMB_1532_ possesses exceptional superparamagnetic properties, with a magnetization saturation of 26.3 emu/g (Figure 1D). Subsequently, we used immunofluorescent (IF) staining to examine whether the Ab-MMB_1532_ were successfully loaded with CD86 antibodies. As shown in Figure 1E, Ab-MMB_1532_ could simultaneously emit both red fluorescence (DiI labeled MB_1532_) and green fluorescence (FITC conjugated CD86) from bubble shells, demonstrating that CD86 antibodies was combined on the surface of Ab-MMB_1532_ through electrostatic adsorption. Besides, the drug loading rate of MB_1532_ and MMB_1532_ were characterized by UV-vis absorption spectroscopy method (Figure S3) ^29^.

### Polarization and identification of M1 macrophages

Macrophages are among the major inflammatory cells in atherosclerotic lesions, especially M1 macrophages. The formation of foam cells by macrophages leads to AS^30^. We have demonstrated that there is high expression of CD86 (a marker of M1 macrophages) within atherosclerotic plaques in apoe^-/-^ mice by IF staining (Figure S4). Therefore, raw264.7 cells were selected for experiments in vitro.

In order to evaluate the phenotypic change of macrophages, IF staining for the M1 macrophage surface marker CD86 was conducted. As shown in Figure 2A, raw264.7 cells underwent a remarkable morphological transformation after treated by LPS for 24 h. Macrophage in control exhibited with rounded and clustered appearance, while LPS group displayed with enlarged cell nuclei and elongated cell shapes, characterized by longer pseudopodia. The degree of cell elongation, which was defined as the longest axis length to the shortest axis length through cell nucleus, was quantified (Figure 2B)^31^. The cell elongation rate of LPS group was about 3 times that of the control group, as well as the average fluorescence intensity of CD86 was approximately 1.3 times that of the controls (*p* < 0.05), confirming that LPS stimulates morphological changes in macrophages, leading to a potential influence on their polarization status (Figure 2C, D). Moreover, the gene expression of M1 macrophages marker including TNF-α and iNOS (Figure 2E-G), as well as the specific proteins iNOS and CD86 (Figure 2H-J), were all remarkably higher than control (*p* < 0.05). In short, the above results indicate that LPS can effectively induce raw264.7 cells to polarize towards the M1 phenotype.

**Fig. 2.**
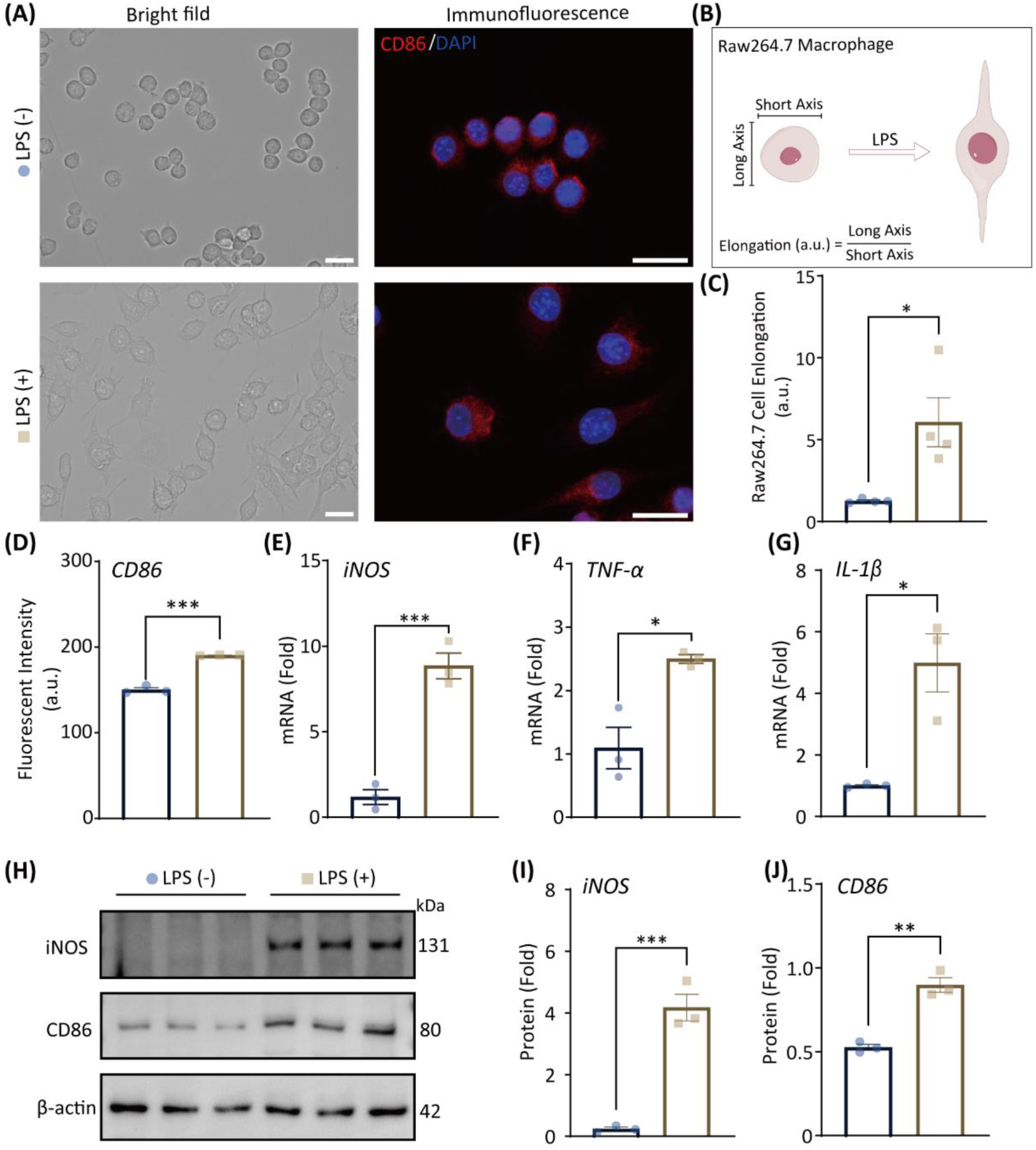
Morphological, mRNA and protein expression changes of raw264.7 macrophages following 24 h of LPS stimulation. (**A**) Representative images of macrophage morphology and CD86 staining. Scale bar: 20 µm. (**B**) Schematic diagram and calculation formula of cell morphology changes, and (**C**) quantitative analysis. (**D**) Semiquantitative analysis of average fluorescence intensity of CD86. (**E-G**) The relative mRNA expression level of iNOS, TNF-α and IL-1β. (H-J) Wb analysis of iNOS and CD86 protein levels. \**p* < 0.05, \*\**p* < 0.01, and \*\*\**p* < 0.001.

BIBR1532 inhibits M1 macrophage proliferation by reducing telomerase activity BIBR1532 has been widely used in research on telomerase function, with reports suggesting its effective inhibition of cell viability in various cancer cell types^23^. However, its utilization in macrophage research remains relatively limited. Therefore, we initially investigated the half-inhibitory concentration (IC_50_) of BIBR on macrophages. Based on the 48 h CCK8 dose-response curve, the IC_50_ value of BIBR1532 for M1 macrophages was calculated as 22.79 μM (Figure 3A). In order to further investigate the potential cell-suppressive effects of BIBR1532 in M1 macrophages, different concentrations of BIBR1532 (5, 10, 20, 30 and 40 μM) were added to M1 macrophages. The results demonstrated that BIBR1532 significantly inhibits cell proliferation in a dose-dependent manner (Figure 3B, C). However, there was no significant difference in cell inhibition rates at drug concentrations of 30 μM and 40 μM.

**Fig. 3.**
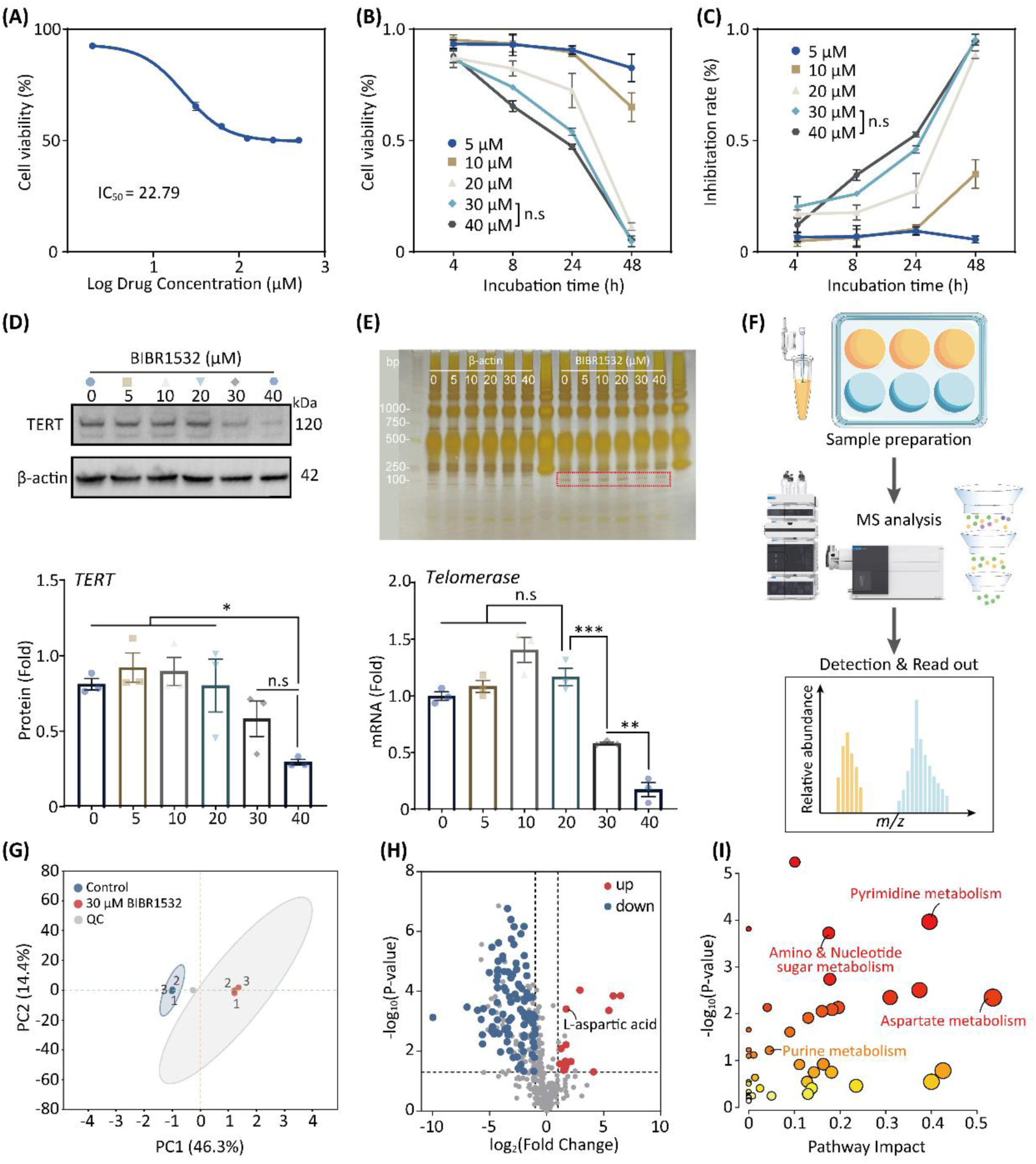
Effect of BIBR1532 on M1 macrophages. (**A**) The IC_50_ value of BIBR1532 was evaluated using CCK8 assays. (**B, C**) Cell viability and inhibition rate were evaluated by CCK8 assay at different time points. (**D**) Wb analysis of TERT protein expression. (**E**) Telomerase activity was detected using a fluorescent RT-qPCR assay kit, and visualizing the results using a protein silver staining kit. (F) Metabolomics sample extraction and analysis schematic diagram. (G) PCA score plot showing differences between the control and the 30μM BIBR1532 groups of M1 macrophages. (H) Volcano plot demonstrating altered metabolite levels between the control and the 30μM BIBR1532 groups. (I) Pathway analysis of significant differential metabolites between the control and the 30μM BIBR1532 groups.

The inhibitory effect of BIBR1532 on cell proliferation is achieved by reducing TERT expression. Western Blot results indicate that TERT protein expression is significantly reduced at a concentration of 40 μM BIBR1532, although there is no significant difference between the 30 μM and 40 μM (Figure 3D). We further investigated the activity of telomerase. As shown in Figure 3E, the telomerase activity assay kit and silver staining results showed a significant decrease of telomerase activity in M1 macrophages at a BIBR1532 concentration of 30 μM, while there was a slight increase at a concentration of 10 μM. We speculated that this might be due to the stress-responsive genes, which were lowly transcribed under normal conditions and robustly induced in response to stress^32^. Similarly, the mRNA expression of TERT provided consistent results, indicating a significant decrease at a BIBR1532 concentration of 30 μM. Following this, the qPCR reaction products were utilized for agarose gel electrophoresis, which clearly indicated a remarkable decrease in TERT gene expression at a BIBR1532 concentration of 30 µM (Figure S5).

In concert, these results concluded that BIBR1532 can inhibit cell proliferation by reducing telomerase activity, especially at the 30 μM concentration. However, the underlying mechanisms require further exploration.

### Metabolomics analysis

The employment of metabolomics has led to new biomarker discoveries and a better mechanistic understanding of diseases with applications in precision medicine^33^. To further investigate the molecular mechanisms behind the decreased telomerase activity and restricted cell proliferation post-BIBR1532 treatment, we utilized ultra-performance liquid chromatography-tandem mass spectrometry (UPLC-MS/MS) to assess the metabolomic characteristics of M1 macrophages treated with and without BIBR1532 (Figure 3F). As shown in the Principal Component Analysis (PCA) score plot (Figure 3G), the two groups exhibit a clear separation trend. After filtering (Fold Change < 0.5 / > 2 and P-value < 0.05), 248metabolites with significant differences were identified in positive ion mode between the control and BIBR1532-treated groups (Figure 3H), and 177 significantly different metabolites in negative ion mode. These differential metabolites were further analyzed using MetaboAnalyst for pathway enrichment. The results indicate that the most affected pathways in both positive and negative ion modes primarily involve pyrimidine & purine metabolism, aspartate metabolism and amino & nucleotide sugar metabolism (Figure 4I). The heatmap of differential metabolites between the control and treatment groups is shown in Figure S6. Notably, compared to the control group, L-glutamine (approximately 0.17-fold) and UMP (approximately 0.13-fold), which are involved in purine and pyrimidine metabolism, were significantly downregulated after BIBR1532 treatment, whereas L-aspartic acid (approximately 3.29-fold), involved in aspartic acid metabolism, was significantly upregulated. Nucleotide levels control telomere lengthening in human cells^34^ and also influence cellular metabolism. After treatment with BIBR1532, M1 macrophages exhibited inhibited telomerase activity, resulting in stagnation of purine, pyrimidine, and nucleoside metabolism. Of note, there was an enhancement in aspartic acid metabolism. Studies indicate that the homeostasis of aspartic acid and asparagine is crucial for cellular fate^35^. We hypothesize that abnormal metabolism of aspartic acid is associated with cell apoptosis.

**Fig. 4.**
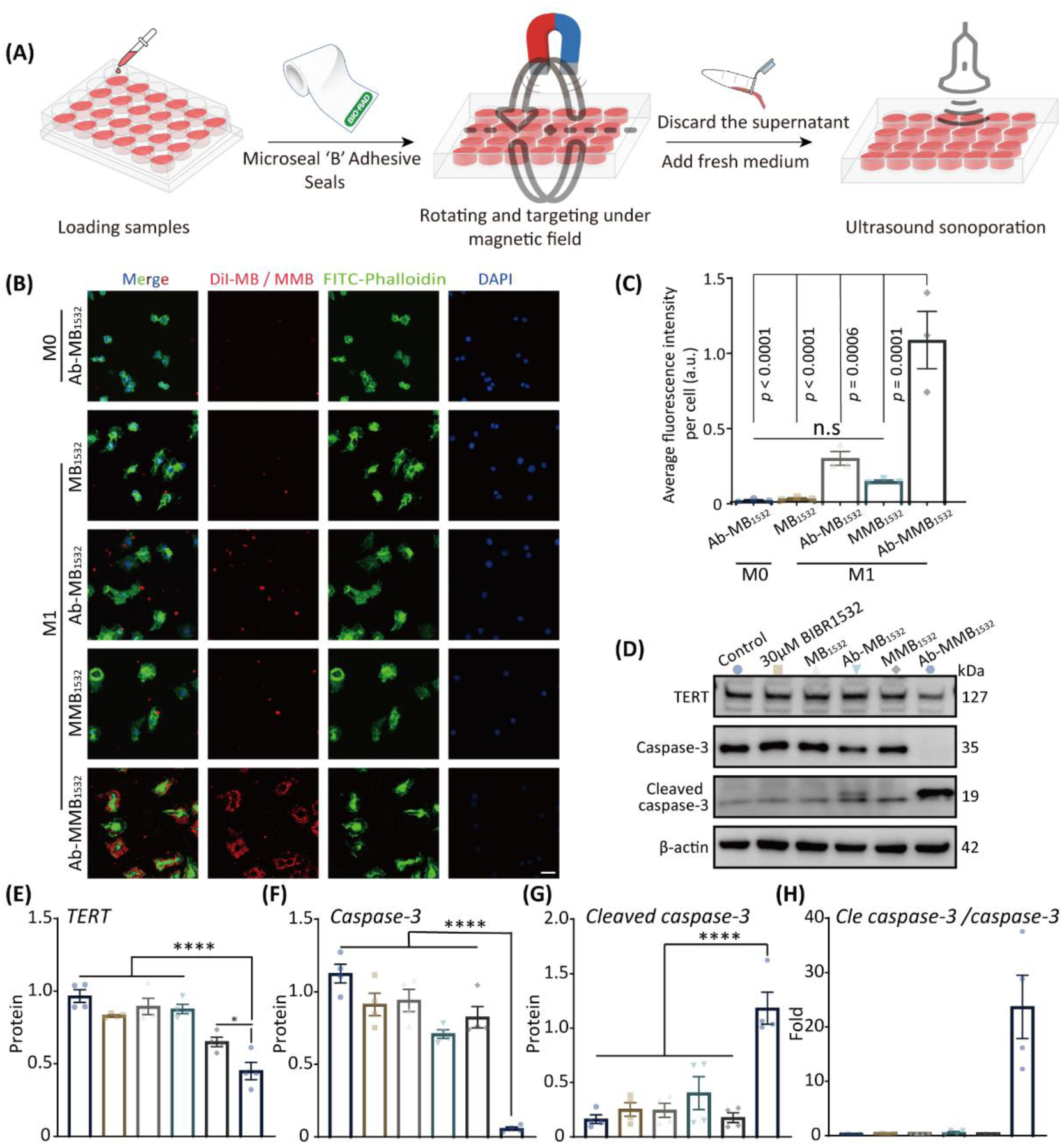
The dual-targeting capabilities and pro-apoptotic potential of Ab-MMB1532. (**A**) Schematic of cell therapy in vitro. (**B**) Assessment Ab-MMB_1532_ targeting specificity by IF. Red represents MB/MMB stained with DiI, green represents FITC-labeled phalloidin staining the macrophage cytoskeleton, and blue represents DAPI staining of cell nuclei. Scale bar: 25μM (**C**) Semiquantitative analysis of average fluorescence intensity. (**D-H**) Western blot analysis of TERT, caspase-3, and cleaved caspase-3. * *p* ˂ 0.05, \*\**p* ˂ 0.01, and *** *p*˂ 0.001.

Therefore, flow cytometry (FC) was used to assess apoptosis in M1 macrophages treated with BIBR1532. FC results demonstrated that the percentage of Annexin-V-positive cells were significantly increased from 5.99% to 9.75% (p < 0.05) after treating M1 macrophages with 10 µM of BIBR1532 for 48 h (5.42% of cells in early apoptotic cells and 4.33% in late apoptotic cells). Furthermore, as the concentration of BIBR1532 increases, the percentage of Annexin-V-positive cells also further increases (Figure S7A). On the other hand, we also investigated the cellular senescence status using a SA-β-Gal staining kit. It was observed that the number of SA-β-Gal-positive cells gradually increased as the concentration of BIBR1532 increased, especially at 30 μM. When the concentration of BIBR1532 reached 40 μM, on the contrary, we did not observe the expression of SA-β-Gal. We speculate that it was due to the toxic effects of BIBR1532, leading to cell death (Figure S7B).

In summary, these results indicate that BIBR1532 suppresses telomerase activity in M1 macrophages, disrupts purine, pyrimidine, nucleoside, and amino acid metabolism, inhibits cell proliferation, upregulates SA-β-Gal expression, thereby promoting cellular senescence and apoptosis.

### Dual-target and pro-apoptotic efficacy of Ab-MMB1532 in vitro

Before assessing the biological effects of Ab-MMB1532, we first evaluated whether the MBs without drugs have any impact on M1 macrophage activity. The CCK-8 assay data reveals that, following co-incubation of M1 with various concentrations of MBs at different time points, M1 activity consistently remains above 90%. This suggests that MBs do not exhibit significant cytotoxicity to M1 macrophages, demonstrating excellent biocompatibility (Figure S8).

Many studies have demonstrated that, compared to single ligands or non-functionalized nanomedicines, delivery systems based on dual ligands are effective^36^. In this study, employed active targeting with CD86 antibodies and magnetic targeting via an external magnetic field, Ab-MMB_1532_ were investigated their biological effects on M1 macrophages in vitro. As shown in Figure 4A, we employed an improved inverted method to simulate the in vitro dual-targeted mediated biological effects, as we described previously^26^.

The finds indicated a substantial concentration of Ab-MMB_1532_ with dual-targeting around M1 macrophages, resulting in a targeting efficiency enhancement of more than 50 % compared with single targeting or non-targeted agents (Figure 4B, C). Additionally, to investigate the specificity targeting of Ab-MB_1532_, we used M0 macrophage as a negative control. Given the nearly absent expression of CD86 in M0, it can be seen from Figure 4B that there was almost no Ab-MB_1532_ around M0. This indicated the specific targeting capability of the Ab-MB_1532_ we prepared. Therefore, dual-target delivery system can greatly improve the specific targeting efficiency to M1 macrophages.

After targeting M1 macrophages, we subsequently applied the ultrasound parameters we determined (Figure S9) for in vitro therapy. The Western blot results demonstrated that the dual-targeting Ab- MMB_1532_ effectively suppress the expression of the TERT protein in M1 macrophages (Figure 4D, E). TERT is a crucial catalytic subunit of telomerase, and its high expression can increase proliferation, anti-apoptosis, and invasion^37^. Therefore, suppressing TERT expression can inhibit the proliferation of M1 macrophages and promote apoptosis. Furthermore, based on the aforementioned metabolomics analysis results, we subsequently examined the expression of apoptosis-related proteins caspase3 and cleaved-caspase3. As anticipated, the results demonstrated that the Ab-MMB_1532_ significantly promote the expression of cleaved caspase-3 while reducing the expression of caspase-3 (Figure 4F, G).

In summary, our data showed that Ab-MMB_1532_ can efficiently and specifically target M1 macrophages. Under the ultrasound cavitation, released BIBR_1532_ can inhibit macrophage proliferation and promote apoptosis by suppressing TERT protein expression, which will contribute to preventing the progression of plaque lesions caused by the activity of M1 macrophages.

### Parallel plate flow chamber assay and targeting capability of MBs in vivo

In order to mimic the magnetic responsiveness and antibody targeting capabilities of MBs in vivo, the targeting efficiency of different microbubble groups was assessed using a custom-designed parallel plate flow chamber under varying shear forces (6 dyn/cm^2^ to 48 dyn/cm^2^) (Figure 5A). As illustrated in Figure 5B, for the 4 types of MBs (MB_1532_, MMB_1532_, Ab-MB_1532_ and Ab-MMB_1532_) investigated, an increase in shear force led to a decrease in the number of captured MBs under the influence of the magnetic field, with the non-targeted group (MB_1532_) exhibiting minimal microbubble capture. Importantly, the dual-targeted group (Ab-MMB_1532_) demonstrated a higher capture percentage even at a high shear force of 48 dyn/cm^2^, outperforming the single-targeted group (MMB_1532_, Ab-MB_1532_). This indicates that dual-targeted MBs exhibit exceptional potential for targeted therapeutic applications, even in high-flow regions such as the aorta.

**Fig. 5.**
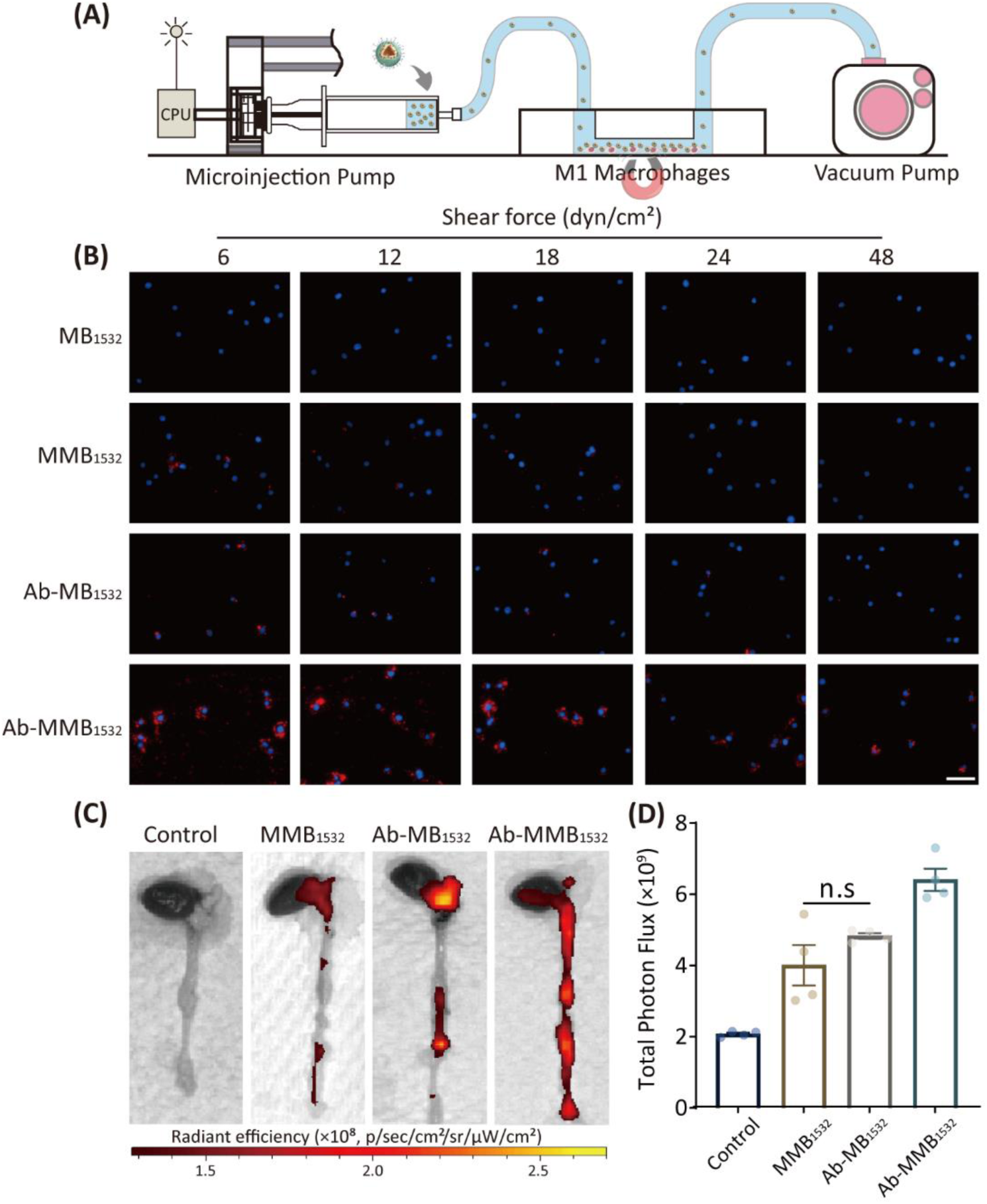
Parallel plate flow chamber assay and targeting AS plaques in apoe^-/-^ mice. (**A**) Schematic of the custom-designed parallel plate flow chamber assay. (**B**) Fluorescence imaging of microbubble capture by M1 macrophages under varying shear forces in the presence of an applied magnetic field (B ≈ 1.2 T). Red represents MBs modified by DiI, and blue represents DAPI. Scale bar: 50μm. (**C**) Representative ex vivo fluorescence images and (**D**) quantitative data of DiD fluorescent signals accumulated in the aorta 24 h post-injection (*n* =4, mean ± SD, *ns*: no significance).

Subsequently, we investigated the targeting capability of *i.v.* administration of MBs on AS plaques in apoe^-/-^ mice. After a 24-hour post-injection period, bright red fluorescence was prominently observed in the aortic arch and abdominal aorta, regions prone to plaque development (Figure 5C). Simultaneously, the Ab-MMB_1532_ administration group exhibited a significantly higher average fluorescence intensity in the aorta compared to the other three groups (*p* < 0.05), with no significant differences observed among the single-targeted groups (MMB_1532_ *v.s* Ab-MB_1532_, *p* > 0.05, Figure 5D). Additionally, fluorescence primarily accumulated in the heart, liver, spleen, and lungs (Figure S10).

### Acute toxicity of Ab-MMB_1532_ in vivo

Prior to evaluating the anti-AS effects of Ab-MMB_1532_ in vivo, we conducted a study on the acute toxicity of Ab-MMB_1532_. After a 4-hour intravenous tail vein injection of Ab-MMB_1532_, the whole blood and major organs of mice were collected, with PBS injection served as a control. Hematological analysis revealed no significant differences in RBC, WBC, PLT, lymphocytes (Lym), neutrophils (GRAN), and hemoglobin (HGB) levels between the Ab-MMB_1532_ and PBS group, and all parameters remained within normal ranges (Figure S11A). H&E staining results demonstrated no apparent pathological changes in major organs such as the hearts, livers, spleens, lungs, and kidneys (Figure S11B). These findings suggest that Ab-MMB_1532_ exhibits good bio-compatibility and holds promise for the treatment of atherosclerosis.

### Therapeutic efficacy on anti-AS

Based on the above promising results, the therapeutic effect of Ab-MMB_1532_ was evaluated in a AS mice model. Following the treatment protocol depicted in Figure 6A, 6-week-old mice were selected and subjected to an 8-week high-fat diet to induce pathological AS. At the end of the treatment period, the entire aorta underwent ORO staining. As shown in Figure 6B, the saline group displayed the highest lesion area as a percentage of the total aortic area, reaching 12.41%. MMB_1532_ and Ab-MB_1532_ treatments both resulted in a moderate reduction in plaque areas by 7.37% and 6.38%, respectively. Notably, the Ab-MMB_1532_ treatment group exhibited a substantial decrease in lesion area as a proportion of the total aortic area, reducing it to 2.35% (Figure 6C). These findings suggest that Ab-MMB_1532_ has a strong anti-AS effect owing to its dual-target properties.

**Fig. 6.**
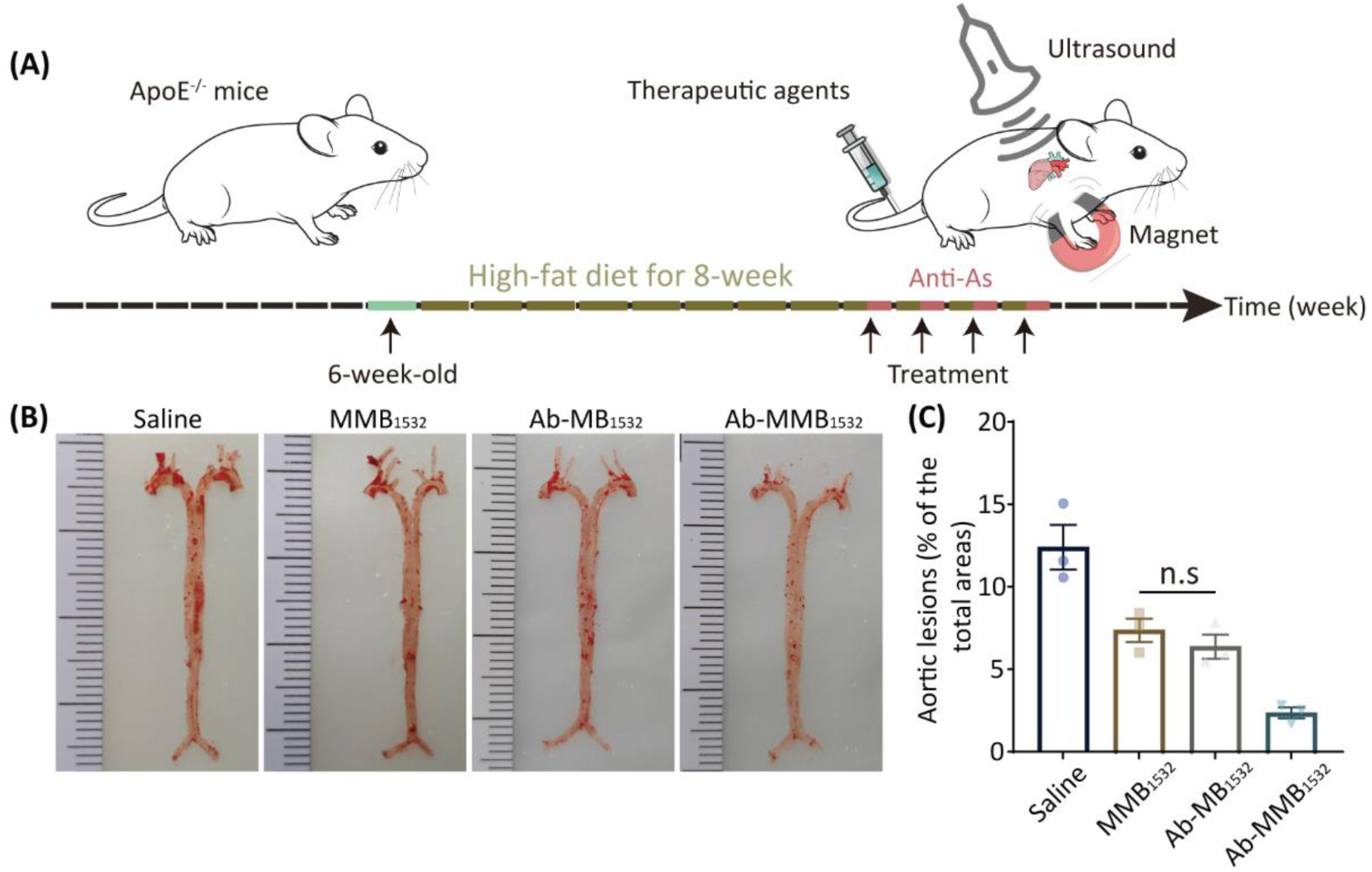
Therapeutic efficacy of AS in apoe^-/-^ mice. (**A**) Schematic representation of the treatment protocol in this study. (**B**) Representative images of aortic ORO staining. (**C**) Quantitative analysis of lesion area (*n* = 3). \**p* < 0.05.

Subsequently, frozen sections of the aortic root were subjected to H&E, ORO, and IHC analysis. As shown in Figure 7A & B, H&E staining of the aortic root in the saline group revealed numerous large plaques and necrotic cores. However, both MMB_1532_ and Ab-MB_1532_ treatment groups exhibited a significant reduction in plaque area (about 0.27 mm^2^ and 0.27 mm^2^, respectively). Notably, the Ab-MMB_1532_ group exhibited the most favorable treatment outcome (0.09 mm^2^).

**Fig. 7.**
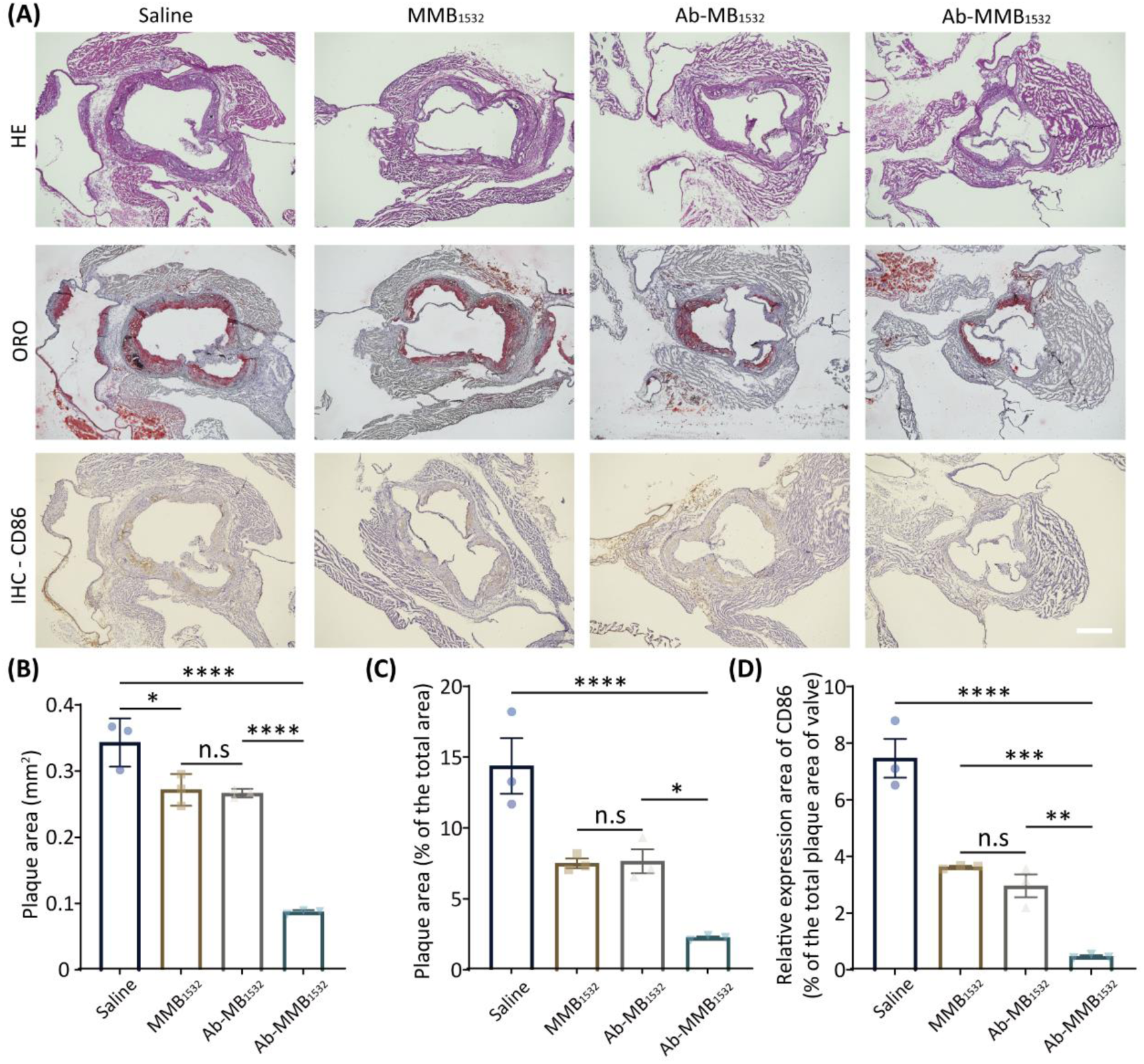
The anti-AS effect of Ab-MMB1532. (**A**) Representative H&E, ORO, and IHC staining (CD86) of atherosclerotic plaque and their quantitative analysis data (**B-D**) (*n* = 3). \**p* < 0.05, \*\**p* < 0.01, \*\*\**p* < 0.001 and *ns*, no significance. Scale bar: 500 µm.

As demonstrated by ORO staining, the therapeutic agents injection group (MMB_1532_, Ab-MB_1532_, and Ab-MMB_1532_) exhibited a reduction in lipid accumulation within AS plaques in treated mice compared to the saline group (Figure 7A). Specifically, the saline group displayed substantial lipid deposition within plaques (approximately 14.4%). In comparison, both MMB_1532_ and Ab-MB_1532_ groups were able to reduce lipid deposition within plaques (7.6% and 7.7%, respectively), with Ab- MMB_1532_ exhibiting the least lipid deposition after treatment (2.3%) (Figure 7C). Furthermore, the analysis of serum LDL, CHO, and TG levels in each groups revealed that the anti-AS effect of the therapeutic agents was not achieved through serum lipid reduction (Figure S12D-F).

It has been reported that M1 macrophages aggravate atherosclerosis progression by stimulating inflammation and lipid engulfment^38^. Therefore, we used IHC to observe whether Ab-MMB_1532_, without affecting serum lipid levels in the precursor, achieves therapeutic goals by regulating M1 macrophages within atherosclerotic plaques. The analysis of CD86, a marker for M1 macrophages, showed a significant decrease in the number of M1 macrophages within the plaques, especially in the group treated with Ab-MMB_1532_. In contrast, the infiltration of M1 macrophages in saline group was about 15 times higher than that in group Ab-MMB_1532_ (*p* < 0.05, Figure 7D). These results suggest that Ab-MMB_1532_ can prevent the infiltration of macrophages and reduce the deposition of lipids, thus achieving the effect of anti-AS.

### Biosafety assessment

After 4-week treatment period, we conducted an assessment of potential side effects associated with various treatment modalities. Our findings revealed that there were no statistically significant differences in hematological parameters (including RBC, Lym, and PLT) and serum lipid level (LDL, CHO, and TG) between MMB_1532_, Ab-MB_1532_, Ab-MMB_1532_, and saline groups (*p* > 0.05, Figure S12A-F). H&E staining results also showed that there were no notable pathological changes in major organs (heart, liver, spleen, lung, kidney) after administrating the aforementioned treatment protocol (Figure S12G). Thus, it further demonstrated that Ab-MMB_1532_ possesses excellent biocompatibility in vivo.

## CONCLUSION

In this study, we developed a dual-target drug delivery system to efficiently target and release drugs at specific sites of AS plaques. In vitro, the Ab-MMB_1532_, which possesses dual-targeting capabilities of CD86 antibodies and magnetic induction, can effectively target to M1 macrophages and release BIBR1532 under sonication, leading to a significant inhibition of M1 macrophage telomerase activity, thereby suppressing cell proliferation and promoting apoptosis of M1. In an AS mouse model, Ab-MMB_1532_ effectively accumulated within AS plaques. Furthermore, in vivo treatment results demonstrated that Ab-MMB_1532_ effectively slow down the advancement of AS following 4 weeks of therapy. Finally, Ab-MMB_1532_ exhibited favorable safety profile without any noticeable adverse effects in mice after long-term administration. Therefore, this novel system with dual-targeting capabilities holds the potential use for the treatment of AS.

## ACKNOWLEDGMENTS

This research was funded by the National Natural Science Foundation of China (Grant No. 82102041); and a Project of Innovation of the Science and Technology Commission of Shenzhen City (JCYJ20210324113804013); Project of International cooperative research of the Science and Technology Commission of Shenzhen City (GJHZ20210705142205017, GJHZ20210705142206019).

## AUTHOR CONTRIBUTIONS

Conceptualization, Hongwen Fei, Jinfeng Xu and Yingying Liu; Data curation, Wei Zeng and Zhengan Huang; Formal analysis, Wei Zeng; Funding acquisition, Jinfeng Xu and Yingying Liu; Investigation, Wei Zeng, Yuanyuan Sheng and Xiaoxuan Lin; Methodology, Yingying Liu; Project administration, Yingying Liu; Resources, Wei Zeng, Zhengan Huang, Yalan Huang, Kaifen Xiong, Xiaofang Zhong, Jiayu Ye, Yanbin Guo and Gulzira Arkin; Software, Wei Zeng and Zhengan Huang; Supervision, Yingying Liu; Validation, Wei Zeng, Jinfeng Xu and Yingying Liu; Visualization, Wei Zeng and Zhengan Huang; Writing-original draft, Wei Zeng; Writing-review & editing, Yingying Liu. All authors have read and agreed to the published version of the manuscript.

